# Isoflurane anesthesia CNS recovery state: ultra-slow oscillations and cellular-stress-responsive gene expression

**DOI:** 10.64898/2026.10.18.760000

**Authors:** Mantre Dehnad, Theresa E. Bjorness, Ashwinikumar Kulkarni, To Thai, Richa Pandey, Genevieve Konopka, Stephan A. Loer, Robert W. Greene

## Abstract

General anesthesia and NREM sleep share slow-wave EEG features, yet whether anesthesia substitutes for physiological sleep remains unclear. We compared recovery from 4h of isoflurane anesthesia (IA) with time-matched sleep and sleep deprivation (SD) in male C57BL/6J mice using continuous EEG/EMG recordings and frontal cortex RNA sequencing. IA recovery produced transient, rebound arousal, reduced early NREM and REM sleep, and, fragmented subsequent NREM bouts, and persistant, ultra-slow EEG-oscillations that normalized within two weeks. Whereas, SD induced consolidated rebound sleep and slow wave activity. Transcriptomic profiling at ZT6 showed that IA and SD induced largely distinct cortical gene-expression programs. IA preferentially enriched integrated stress response and unfolded protein response genes, whereas SD showed a comparatively orthogonal, growth-permissive, mTOR-associated profile. Although IA-induced gene-expression suggests cellular stress, corticosterone measurements argued against systemic stress as the primary driver. These findings indicate that isoflurane recovery is neurophysiologically and molecularly distinct from restorative sleep.

## Introduction

General anesthesia is one of the most widely applied interventions in medicine, enabling hundreds of millions of surgeries each year and transforming formerly impossible surgical operations into routine care^1,2^. Understanding how anesthetic drugs alter neural activity and recovery is clinically important for optimizing perioperative brain health, postoperative cognition, and the management of prolonged or repeated anesthetic exposures as the drugs clearly interfere with physiological neuronal function in order to impose an anesthetic state^3,4^. The anesthetic state shares several features with the natural state of NREM sleep, including reduced responsiveness to external stimuli, loss of consciousness and stereotyped electroencephalographic (EEG) dynamics^5^. However accumulating evidence indicates that anesthesia- and sleep-associated oscillations arise from distinct circuit mechanisms and exhibit different patterns of connectivity and thalamocortical coordination, consistent with their being distinct neurophysiological states^6,7^.

In this study we focus on the volatile anesthetic isoflurane, which, along with other volatile anesthesics such as sevoflurane and desflurane, is widely used for maintaining general anesthesia during procedures lasting hours, including major abdominal, thoracic, and neurosurgical operations, as opposed to intravenous anesthetics such as propofol which are generally reserved for induction (particularly in the United States) or shorter procedures (lasting shorter than an hour), with agent selection shaped by procedure and patient factors.

These volatile agents have long been described as inducing a state resembling non-rapid eye movement (NREM) slow-wave sleep (SWS), because of striking EEG similarities, including prominent delta activity^5,8^. Under their influence, involving potentiation of GABA_A_-receptor-mediated inhibition and activation of potassium channels, cortical and thalamic networks generate high-amplitude slow wave activity (SWA), also known as delta oscillations (∼0.1-4 Hz), and synchronized slow rhythms that visually mirror those seen in physiological NREM sleep^2,8^. This resemblance, first recognized decades ago and confirmed by contemporary recordings, has led to the prevailing notion that anesthesia recapitulates elements of sleep, particularly the deep-sleep component.

Yet a closer look reveals perplexing differences. Although the delta power observed during anesthesia may appear indistinguishable from that of slow-wave sleep, post-anesthetic sleep is disrupted with evidence of altered sleep architecture including fragmented or less consolidated deep sleep stages, and changes in REM duration^9–12^. In addition, reproducible spectral distinctions, including prominent enhancement in the ultra-slow (<1 Hz) oscillation range during and immediately following anesthestia, supports the idea that anesthetic slow rhythms are not equivalent to physiological sleep slow rhythms^6,13–15^. Whether such ultra-slow activity persists into recovery and generalizes across arousal states, or associates with chages in cortical cellular state has remained unclear. These differences raise the possibility that time spent under anesthesia may not substitute for physiological NREM.

Prior work suggests the extent to which anesthesia “counts” toward sleep recovery varies by anesthetic agent and sleep domain. Anesthesia-associated slow waves can blunt the expected slow-wave activity (SWA) rebound after sleep deprivation, that could be interpreted as a reduction of specific NREM pressure signals, often considered as a reflection of reduced sleep need^16^. However, there are examples of dissociation of SWA rebound from sleep need, including, blunting of rebound SWA resulting from increased arousal^17,18^ or from reduced adenosine activation of ADORA1s^19^ independently of sleep need.

Here, we examine how a clinically relevant exposure, four hours of isoflurane anesthesia, affects both the neurophysiologic and molecular recovery trajectory of the cortex. We used dense 48-hour EEG/EMG recordings before and after anesthesia, compared with time-matched sleep deprivation (SD), to characterize the time course of post-isoflurane changes in sleep architecture and cortical activity. We then probed the molecular activity of this altered state using bulk RNA sequencing to ask whether the pattern of genomic expression might be similar but opposite in polarity to that in response to SD as might be expected in response to an experimentally induced increase of sleep activity and function. These studies provide evidence that, rather than serving as a physiologic substitute for sleep, volatile anesthesia elicits a distinct recovery state with its own neurophysiologic signature and gene-expression signature.

## Results

### Arousal and sleep architectures are altered after isoflurane exposure

The responses to sustained general anesthesia with respect to arousal, EEG and state architecture remain unresolved. To address this, we compared the response that follows 4 h of undisturbed spontaneous sleep to the response following 4 h of anesthesia induced, sleep-like state, reasoning that the latter response, if functionally sleep-like, should resemble the former with respect to these features. We recorded 48 h of baseline sleep (control sleep, CS), administered either 4 h of isoflurane anesthesia (IA; *n* = 7) at lights-on (ZT0) or 4 h of SD (*n* = 8), and then, monitored 48 h of post-anesthetic recovery (Fig. 1A). Relative to the corresponding baseline hour (after four hours of ad lib sleep), the first hour of recovery from anesthesia revealed a striking divergence in arousal-state. IA animals remained predominantly awake, with markedly reduced NREM and REM sleep, consistent with an arousal-biased recovery(Fig. 1B,C; REM in Fig. S1A, Table S1). Wake time during the first recovery hour was increased in IA relative to baseline (mean ± SEM: 56.57 ± 1.46 min vs 22.88 ± 3.16 min; *p* = 0.064), while NREM and REM were correspondingly reduced (NREM: 2.91 ± 1.16 min vs 34.57 ± 2.97 min; *p* = 0.015; REM: 0.10 ± 0.10 min vs 2.45 ± 0.56 min; *p* = 0.40) (Tables S1).

**Figure 1.**
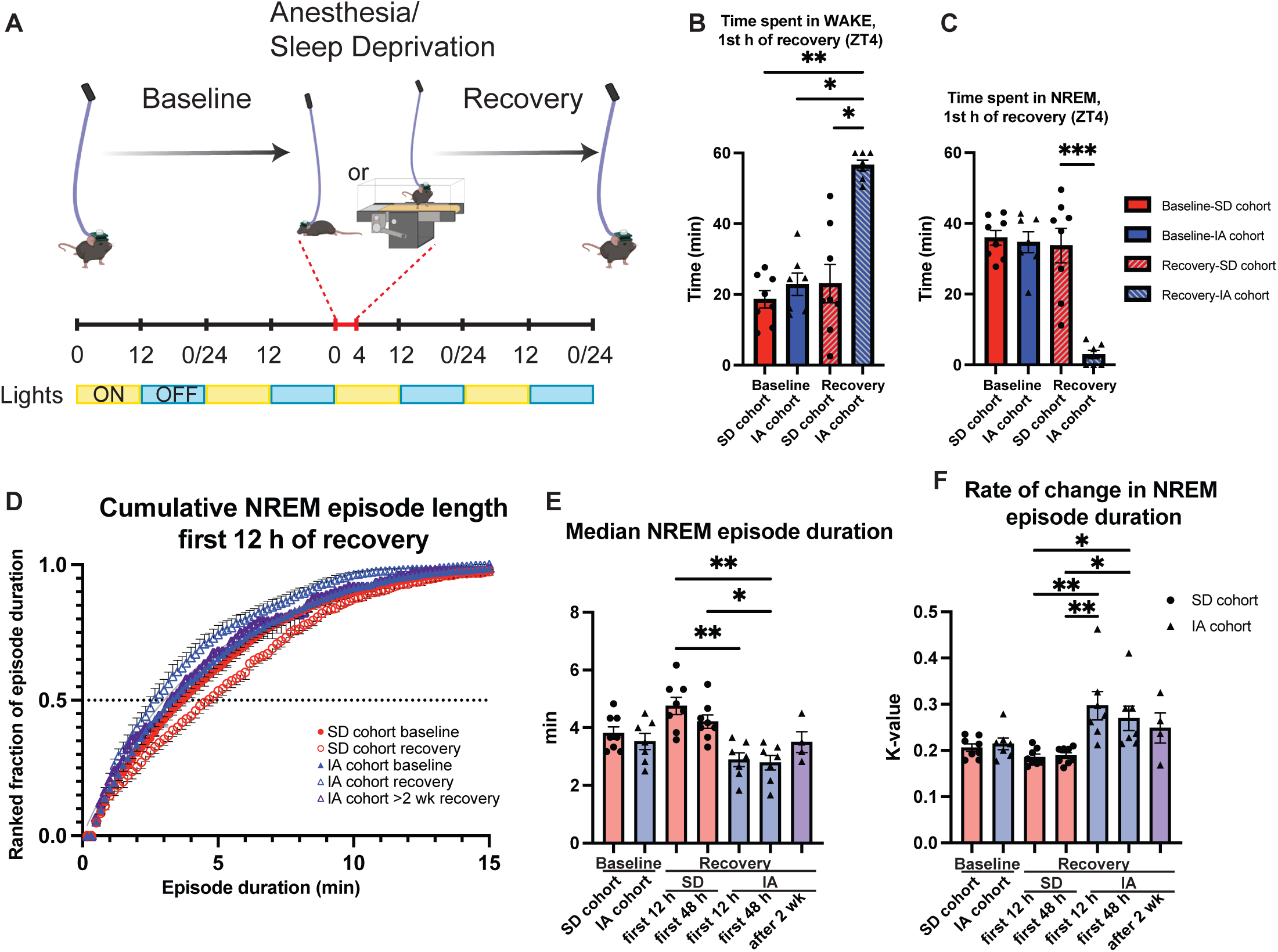
Isoflurane disrupts early recovery sleep architecture and fragments NREM bouts. **(A)** Experimental paradigm: 48 h baseline recording, followed by 4 h of isoflurane anesthesia (IA; *n* = 7) or sleep deprivation (SD; *n* = 8) at lights-on (ZT0), and 48 h of recovery monitoring. **(B,C)** Time spent in wake (B) and NREM sleep (C) during the first recovery hour (ZT4), expressed relative to the corresponding baseline hour. **(D)** Ranked cumulative distributions of NREM episode lengths during the first 12 h of recovery (IA, blue triangles; SD, red circles). For reference, IA NREM episode distributions normalized to baseline after two weeks of recovery are shown (purple triangles). **(E,F)** Quantification of NREM sleep architecture, showing median NREM episode duration (E) and the rate constant **K** (F) derived from fits to cumulative episode-length distributions. Cumulative NREM episode-length distributions were fit using a single-exponential plateau model Y=Y_M_ -(Y_M_-Y_0_)*exp(-k*x) implemented in GraphPad Prism using least-squares regression without weighting (Y_0_ = 0, Y_M_ = 1). The parameter **K** reflects the rate of decay of the distribution, with larger values indicating a higher proportion of short NREM episodes (i.e., greater fragmentation). At baseline, IA and SD cohorts did not differ in median NREM episode duration or **K**. During early recovery (first 12 h), IA exhibited significantly shorter NREM episodes and higher **K** values compared to SD (*p*adj < 0.01 for both metrics). Group comparisons were performed using a Kruskal-Wallis test with Dunn’s post hoc correction. Data are shown as mean ± SEM unless otherwise indicated. * *p*adj < 0.05, ** *p*adj < 0.01, *** *p*adj < 0.001, **** *p*adj < 0.0001.

The marked, response differences, might reflect a kind of pharmacological sleep-loss, so responses to SD with the same duration and circadian timing were examined. In contrast to CS or IA, SD animals rapidly re-entered a sleep-rich state characteristic of homeostatic rebound. These early differences indicate that time spent under anesthesia is not equivalent to either physiological sleep or to restorative rebound sleep after SD. Whereas recovery from SD is characterized by a rapid transition into consolidated sleep, emergence from isoflurane instead produces a transient period of hyper-arousal followed by gradual normalization toward baseline wakefulness. Furthermore, significantly increased sleep, induced by DREADD activation of VLPO sleep center neurons, elicits a non-significant sleep EEG response, unlike IA^20,21^. This divergence confirms prior work showing that anesthesia and sleep deprivation engage distinct, recovery dynamics despite superficial similarities during unconsciousness^6,11,22–26^.

### Post-anesthetic NREM sleep is fragmented

We next examined whether these state-level responses reflected altered sleep architecture. At baseline, IA and SD cohorts exhibited similar NREM episode durations and comparable rate constants (K ≈ 0.20; Fig. 1E,F). Consistent with prior reports of post-anesthetic sleep fragmentation^9–12^, following IA, NREM episodes became significantly shorter and more fragmented relative to baseline, reflected by reduced median episode duration and increased K values. In contrast, SD animals showed modest changes in NREM consolidation that were consistent with rebound sleep and did not exceed baseline variability (Fig. 1D-F, Fig S2). Sleep fragmentation is widely recognized as a hallmark of poor sleep quality and, in many contexts, a signature of pathological sleep.

### Ultra-slow (0-1 Hz) oscillations dominate post-anesthetic EEG

Because both sleep deprivation and anesthesia are associated with prominent slow oscillations with greater energy than normal sleep, we next examined how cortical spectral dynamics diverged during recovery from each state. At baseline, EEG power spectra were broadly similar across cohorts in wake, NREM, and REM sleep (Fig. 2A,D,G).

**Figure 2.**
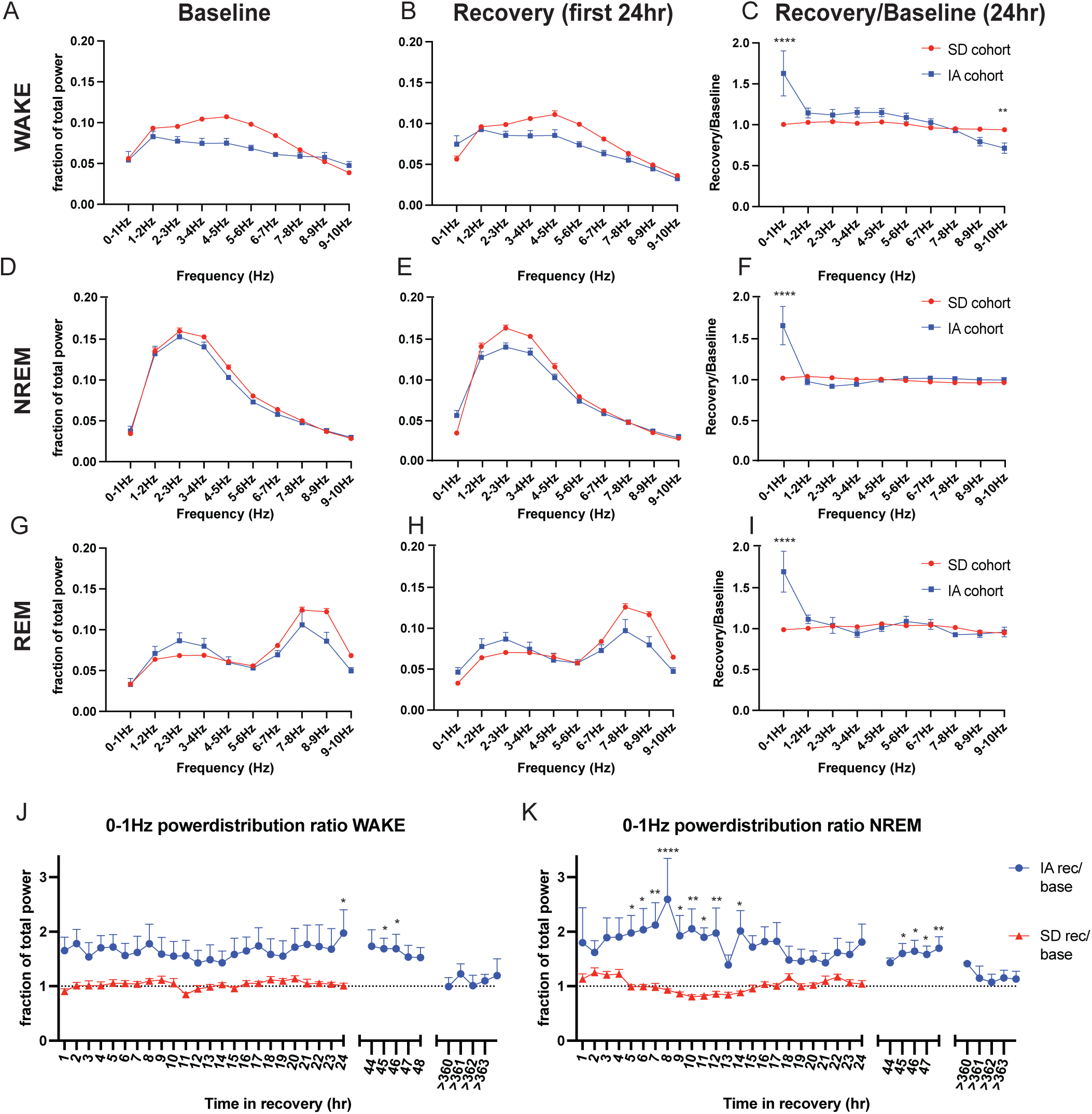
Isoflurane induces a persistent ultra-slow oscillatory bias during recovery. **(A-I)** EEG power distributions (0-10 Hz) during wake (A-C), NREM (D-F), and REM (G-I), comparing baseline control sleep (CS) with the first 24 h of recovery after IA or SD. **(A,D,G)** Baseline fractions of total power within each vigilance state. (B,E,H) Power distributions during the first 24 h of recovery. **(C,F,I)** Recovery-to-baseline power ratios for each frequency bin. **(J,K)** Recovery-to-baseline ratios for ultra-slow (0-1 Hz) power during wake (J) and NREM (K), plotted over two weeks following IA and over the first 24 h following SD. Group differences in recovery-to-baseline power ratios (C,F,I) were assessed using a two-way ANOVA with factors of group (IA vs SD) and frequency bin, including a group x frequency interaction. Group differences in ultra-slow (0-1 Hz) power over time (J,K) were assessed using a two-way ANOVA with factors of group (IA or SD) and time. Post hoc comparisons were corrected using the Šídák method. The largest IA–SD divergence occurred in the ultra-slow (0-1 Hz) band (C,F,I). Data are shown as mean ± SEM. * *p*adj < 0.05, ** *p*adj < 0.01, *** *p*adj < 0.001, **** *p*adj < 0.0001.

During the first 24 h of recovery (Fig. 2B,E,H), IA mice exhibited a pronounced and pervasive increase in ultra-slow (0-1 Hz) power across all arousal states, whereas SD mice showed relatively greater power in higher low-frequency bands corresponding to their baseline spectral profile. Specifically, in wake, SD mice maintained modestly higher power in the 2-7 Hz (∼3% mean increase relative to baseline, variability shown as mean ± SEM in Fig. 2B), and in NREM, higher power in the 2-5 Hz range (∼3% increase, Fig. 2E). In contrast, IA animals showed a marked redistribution of spectral power toward the 0-1 Hz band. Normalizing recovery spectra to baseline underscored this divergence, revealing a ≥50% elevation in 0-1 Hz power after IA across wake, NREM, and REM sleep (Fig. 2C,F,I). This ultra-slow power enhancement persisted for more than 48 h and, finally, returned to baseline when tested after two weeks (Fig. 2J,K; REM in Fig. S1B).

The 0-1 Hz band encompasses the canonical cortical slow oscillation classically associated with NREM sleep^13,27^. Notably, however, unlike physiological NREM slow oscillations, the IA-associated enhancement was not state-specific, as it was observed during wakefulness and REM sleep along with NREM. Consistent with these observations, two-way ANOVA revealed a significant treatment x frequency interaction (Fig. 2C,F,I), confirming its association with IA. Such state-independent ultra-slow wave activity is inconsistent with physiological sleep architecture and has been previously linked to altered synaptic integration and impaired information processing^15,28^.

Both the fragmented NREM pattern following IA and the emergence of significantly increased state-independent, low frequency power indicate an atypical recovery state, one that stands in stark contrast to either sleep following physiological sleep or that of post-SD rebound sleep. This motivated us to ask whether this aberrant recovery state has a functional correlate at the molecular level in the forebrain.

### Isoflurane anesthesia elicits distinct cortical transcriptional signatures

To better understand the functional implications of the electrophysiological and sleep-architectural divergence between IA and physiological sleep, we characterized differences in protein encoding mRNA using bulk RNA-seq in the frontal cortex (FC). FC mRNA was collected two hours after IA recovery and analyzed for differential gene expression relative to an ad libitum control sleep cohort (CS) at the same circadian time during the light phase (ZT6). All libraries (n=26) from CS (n=13 total; 6 CS matched to IA cohort, compared to 7 CS matched to SD cohort from published dataset^29^), IA (n=6), and SD (n=7) cohorts were analyzed jointly. Only reads of genes shared in both cohorts (8420 genes) were normalized by RPKM and covariants (batch, RIN value, hemisphere) were regressed out before differential expression analyses (supplemental table S2).

The CSxIA comparison produced 472 DEGs (FDR < 0.05, |log2FC| ≥ 0.3; 305 up, 167 down; supplemental table S2; including robust induction of immediate-early genes *Fos*, *Egr2*, and *Arc* (Fig. 3A, B). Notably, the IA DEGs were significantly enriched for genes associated with cellular stress response as described below (113 stress DEGs/ 359 DEGs; supplemental table S4-S5).

**Figure 3.**
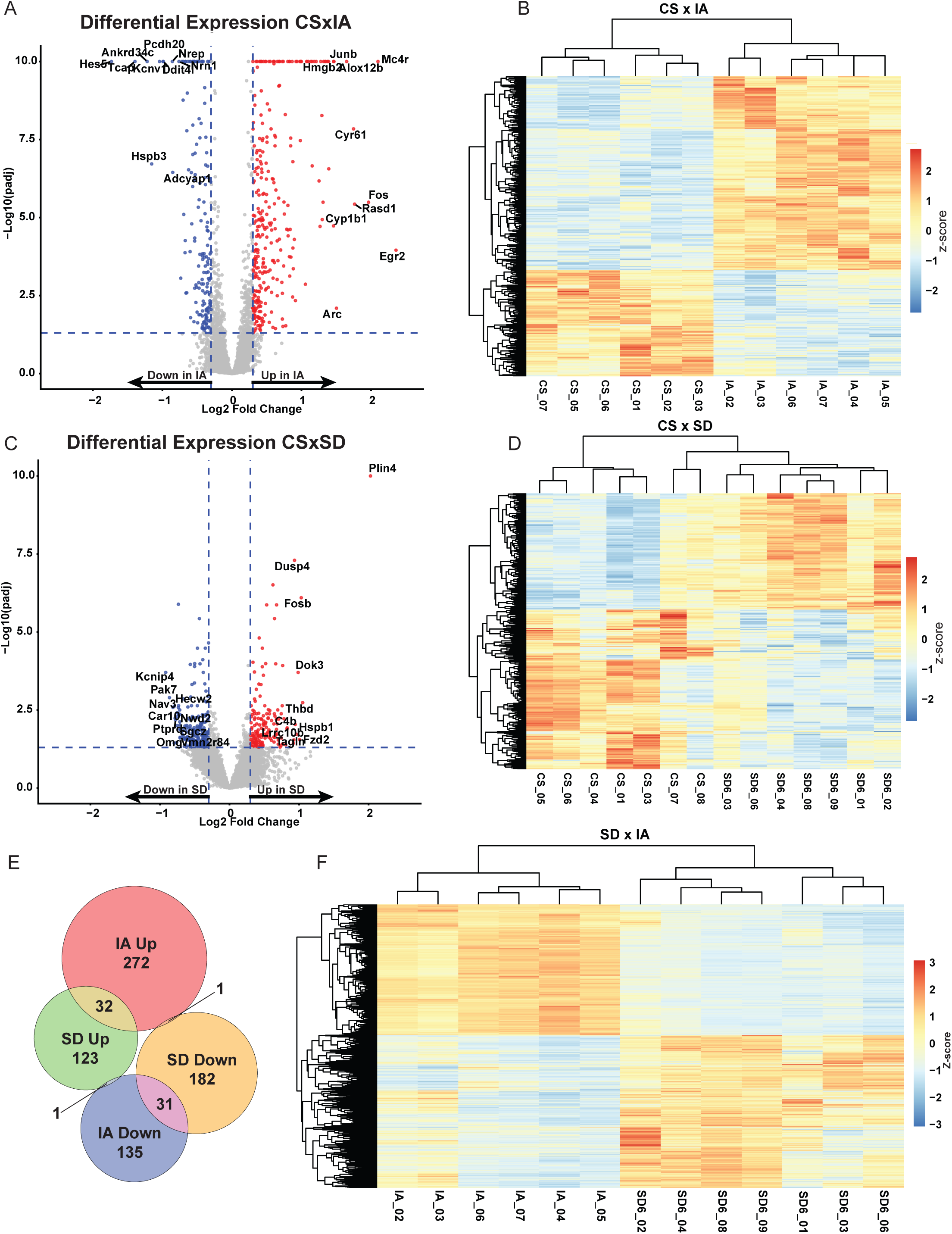
Isoflurane and sleep deprivation elicit distinct cortical transcriptional signatures. **(A)** Volcano plot of differential gene expression for IA vs CS (305 upregulated, 167 downregulated genes). (B) Heatmap of *z*-scored expression for the 472 IA-regulated genes. (C) Volcano plot of differential gene expression for SD vs CS (156 upregulated, 214 downregulated genes). (D) Heatmap of *z*-scored expression for the 370 SD-regulated genes. (E) Venn diagram showing overlap of differentially expressed genes across IA and SD (IA_up ∩ SD_down = 1; IA_up ∩ SD_up = 32; IA_down ∩ SD_up = 1; IA_down ∩ SD_down = 31). (F) Heatmap of *z*-scored expression of 642 genes differentially regulated across IA and SD conditions.

### The CSxSD DEG set is distinct from the CSxIA DEG set

Since the molecular fingerprint of IA has marked dissimilarities from CS, suggesting functional dissimilarity from sleep, we considered whether it may have similarities with SD that could occur if IA engages a molecular program resembling the response to sleep loss, but perhaps opposite in polarity as might be consistent with anesthesia-enhanced sleep. Using bulk RNA-seq collected from the FC of another two cohorts (n=14, n=7 for SD and matched CS) that underwent a validated 6hr SD protocol (published dataset^29^) at matched circadian time and considering only those genes expressed in both of the two paired cohorts, CSxSD yielded 370 DEGs (156 up, 214 down) with prominent up-regulation of *Fosb*, *Plin4*, and *Dusp4* (Fig. 3C,D; supplemental table S4). Comparing the CSxSD DEGs to CSxIA DEGs, there is greater sample to sample variability in the CSxSD set (supplementary table S4). Many of the DEGs are significantly affected by only one of the two sets of comparisons between the two paired cohorts. Most of the DEGs showed opposite polarity of change in expression as illustrated by the heat map (Fig. 3F) and as might be expected if IA promotes a gain in sleep function (Fig. 3E). However, only 65 genes overlapped (were the same) between IA and SD (65/777; ∼10% of the union; Fig. 3E,F,) andnly 2 of these exhibits opposite polarity (significantly upregulated in one comparison and downregulated in the other; Fig. 3F), inconsistent with an IA gain of sleep function.

Together, these findings demonstrate that IA and SD engage fundamentally different cortical transcriptional expression changes from CS, consistent with divergent mechanisms of network recovery and plasticity after anesthesia versus physiological sleep loss.

### IA induces an intracellular stress response-enriched gene set

Our group’s prior work has shown that the molecular response to sleep loss is a sensitive indicator of the brain response to extended wakefulness, with specific synaptic, and metabolic programs predicting the trajectory of rebound sleep and cortical restoration^29,30^. Within this framework, we asked whether the ontology of DEGs might provide some insight into the cellular state of frontal brain tissue, associated with the abnormal post-IA electrophysiology and sleep-state architecture.

We curated a composite stress-gene set comprising integrated stress response (ISR) and unfolded protein response (UPR) genes from Gene Ontology annotations (https://www.informatics.jax.org/vocab/gene_ontology/) and retained only those expressed in our FC samples (Supplementary Table S3). We found significant enrichment of the curated stress genes in IAxCS DEGs (113/472 stress DEGs; Fisher’s Exact test two-tailed P value <0.0001; Supplementary Table S4). In contrast, there were relatively few stress genes significantly affected by SD (61/370; Fisher’s Exact test two-tailed P value 1.000, Supplementary Table S5). A polarity specific Venn diagram of the numbers and overlap of the stress DEGs for IA and SD shows an overlap of only 17/169 stress DEGs (Fig 4A). When considering all stress genes expressed in the IA and SD samples, polarity of expression was generally opposite between the two conditions as illustrated by a heat map of stress genes (Fig. 4C).

**Figure 4.**
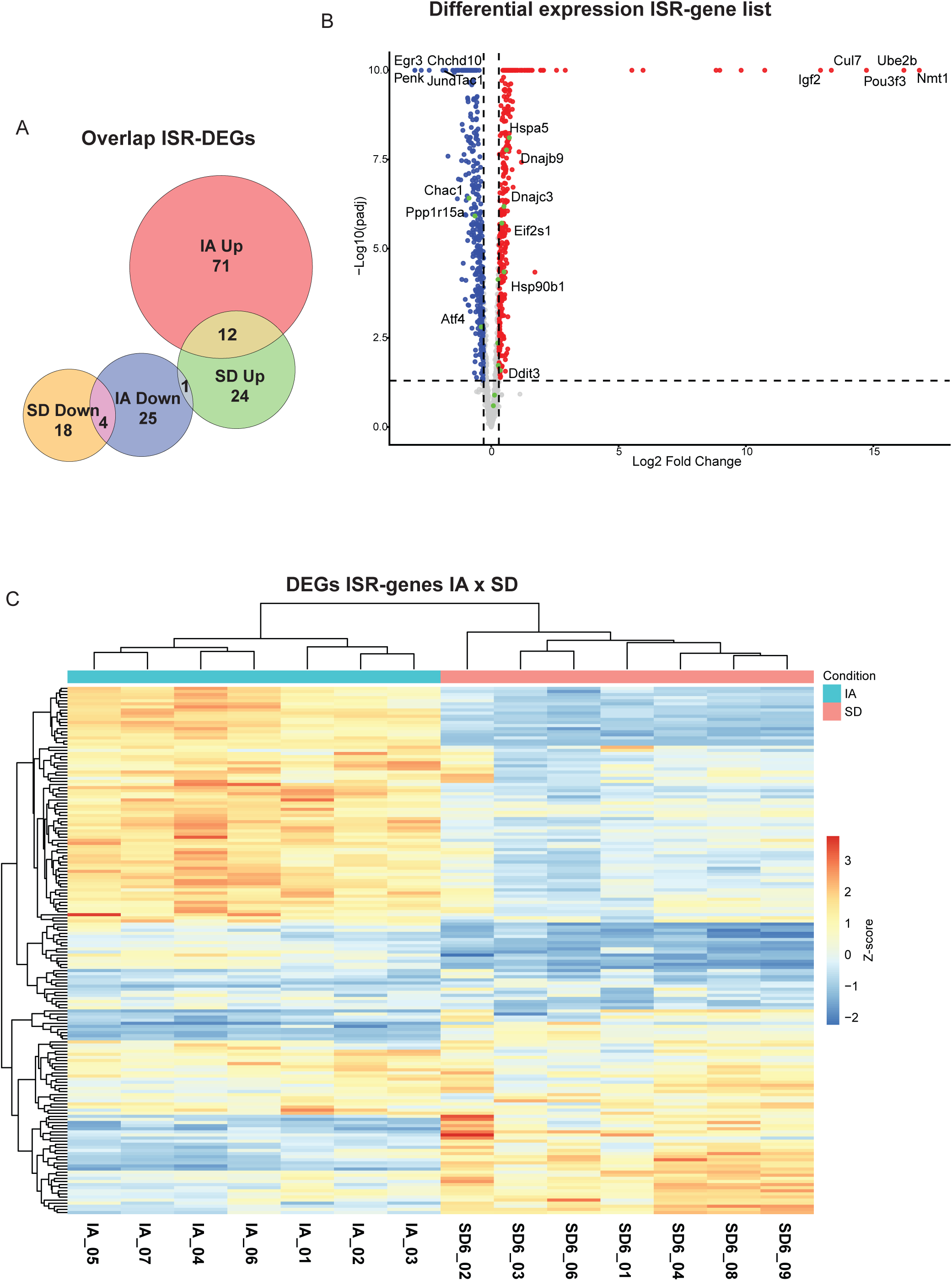
An ISR/UPR-enriched stress-gene program distinguishes anesthesia from sleep deprivation. **(A)** Venn diagram summarizing enrichment of a curated ISR/UPR/cellular stress gene set (Supplementary Table S4-S5) across IA and SD conditions (counts as reported in the main text). (B) Volcano plot of IA vs SD differential expression restricted to the stress-gene universe, with key UPR pathway genes highlighted. (C) Heatmap of significantly differentially expressed stress-related genes demonstrating opposite polarity between anesthetic and sleep-deprivation conditions.

Canonical ISR/UPR markers, including *DDIT3/CHOP*, *PPP1R15A/GADD34*, *EIF2AK3/PERK*, and *XBP1*, were robustly elevated after IA (Fig. 5A) indicating engagement of PERK-eIF2α signaling and adaptive proteostatic remodeling^31,32^. This suggests IA induces a state of cellular stress in association with enhanced state-independent ultra-slow oscillations.

**Figure 5.**
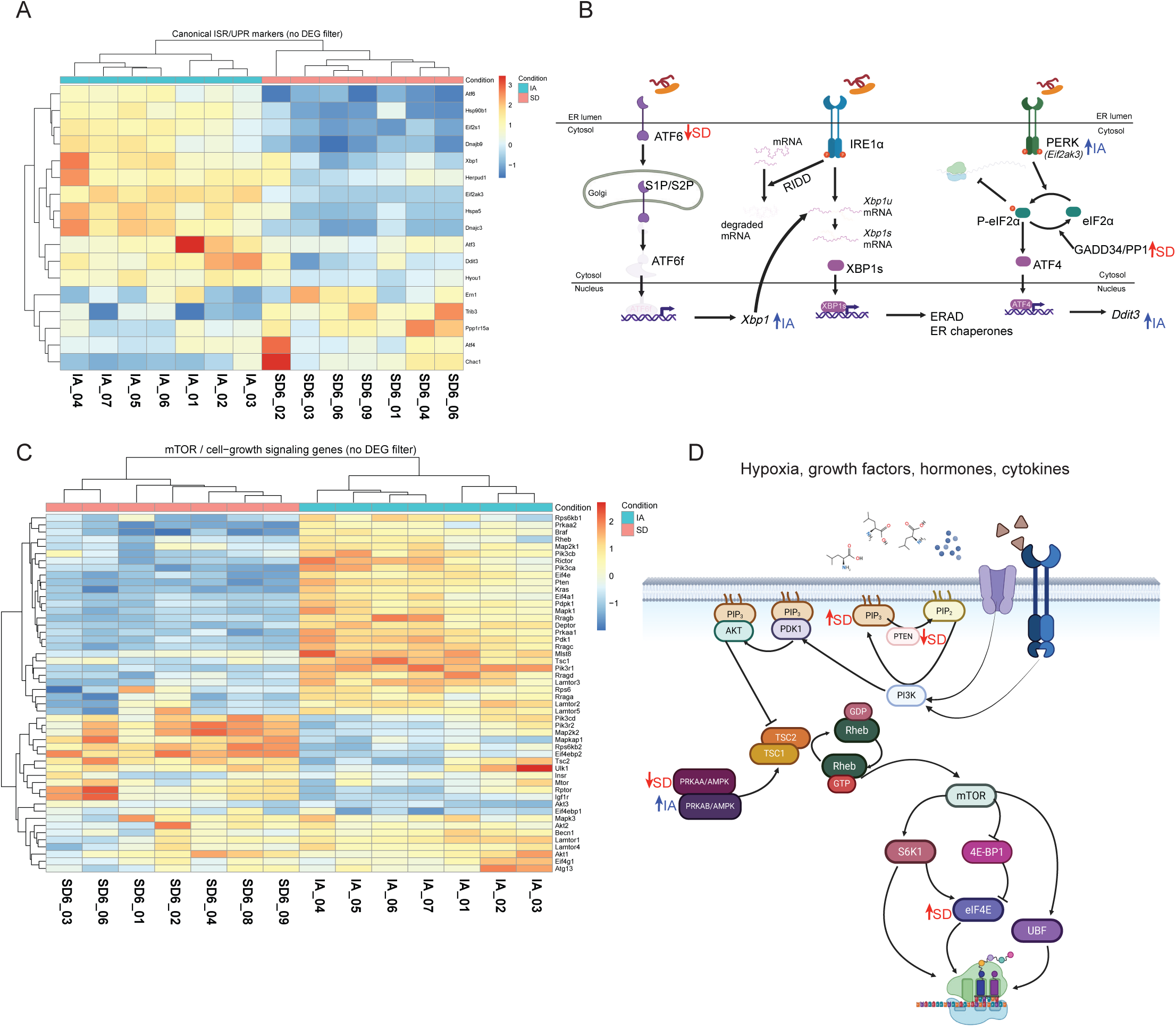
Divergent regulation of UPR and mTOR signaling pathways following isoflurane anesthesia and sleep deprivation. **(A)** Heatmap of *z*-scored expression for canonical unfolded protein response (UPR) genes, comparing IA and SD relative to control sleep. Genes show opposite regulation across conditions, with ISR/UPR components elevated after IA and reduced or unchanged after SD. (B) Schematic overview of the three major branches of the UPR (PERK–eIF2α, IRE1–XBP1, and ATF6). Genes significantly differentially expressed in either IA or SD are highlighted, illustrating preferential engagement of PERK/IRE1 pathways following anesthesia. (C) Heatmap of *z*-scored expression for genes within the mTOR signaling pathway, demonstrating a mirrored expression pattern between IA and SD, with relative suppression following IA and permissive/anabolic regulation following SD. (D) Schematic of the mTOR signaling pathway with differentially expressed genes highlighted, emphasizing divergent regulation of growth- and metabolism-associated nodes between anesthetic recovery and sleep-deprivation rebound. Heatmaps display genes significantly differentially expressed in at least one condition (FDR < 0.05). Color scale represents relative expression (*z*-score) across samples.

An assessment of key metabolism-related genes indicates CSxSD DEGs included down-regulation of ATF6-pathway components and growth-restricting nodes such as *PTEN* and *PRKAA* family members, consistent with transient anabolic mTORC1 activation during early rebound, restorative sleep function. This contrasts with the up-regulation of canonical ISR/UPR DEGs induced by IA that likely depressed the anabolic bias, associated with SD DEG responses (Fig. 5C).

Together, the opposite transcriptional, protein-metabolism-related expression patterns indicate that anesthesia and sleep deprivation drive fundamentally distinct molecular recovery programs, one stress-adaptive (IA), the other growth-permissive (SD), underscoring that anesthesia represents a biologically unique cortical state rather than a molecular substitute for restorative sleep.

### Glucocorticoid profiling rules out systemic stress as a confound

Because glucocorticoids can influence both sleep architecture and gene expression, we measured plasma corticosterone at ZT6 across SD, IA, and a separate restraint-stress positive control. Corticosterone levels after IA and SD remained within the expected circadian range^33–36^ and were not elevated above baseline, whereas restraint stress produced the anticipated increase (Fig. S3). These results are consistent with prior work showing that SD engages the hypothalamic-pituitary-adrenal (HPA) axis only mildly and transiently, and argue against a glucocorticoid-mediated explanation for the ISR/UPR-rich transcriptional program observed after IA^37^.

## Discussion

Here, by pairing continuous EEG/EMG recordings with circadian phase-matched cortical transcriptomics, we characterize the brain state following a clinically relevant 4-hour exposure to isoflurane and compare it directly with the homeostatic response to SD. Together, our data show that IA produces a recovery state that is distinct from both physiological sleep and recovery from sleep loss. IA induces a rebound arousal, increases synchronized ultra-slow oscillatory activity, fragments early NREM sleep, and engages a distinct cellular stress-adaptive molecular program that diverges orthogonally from both physiological sleep and from the response to SD^29^.

Thus, anesthesia, in contrast to physiological sleep, appears to perturb the ensuing level of arousal and sleep architecture^6,8,24^. This pattern aligns with reports of sleep fragmentation and altered circadian dynamics in patients during the first nights after surgery^38–40^, although postoperative sleep disruption is influenced by additional factors including pain, post-operative medications, and the hospital envirntmont. Importantly, the electrophysiological abnormalities following IA persisted well beyond behavioral emergence. The distinction between increased sleep and the post-anesthestic EEG phenotype is further seported by experiments in which activation of neurons in the ventrolateral preoptic area (VLPO) parkedly increased NREM sleep without altering the NREM EEG spectrum^21,22^. Thus, simply increasing physiological sleep is insufficient to reproduce the spectral changes observed following IA.

Transcriptomic analyses indicated that isoflurane robustly up-regulated canonical ISR/UPR effectors, including DDIT3/CHOP, PPP1R15A/GADD34, EIF2AK3/PERK, and XBP1, hallmarks of transient translational attenuation and adaptive proteostasis^31,32,41,42^. The molecular transcriptional response to sleep loss reflects cellular changes that support an anabolically biased, restorative function of sleep^29^. In this categorical respect, SD produced an oppositely polarized signature from that of IA, characterized by relative down-regulation of ATF6-pathway components and negative regulators of mTOR signaling (PTEN, PRKAA family), consistent with rebound-associated restorative programs^43–45^. Although SD’s impact on mTOR signaling can vary by brain region and experimental paradigm, the FC-restricted, ZT6-matched design here reveals an early cortical dichotomy: anesthesia favors ISR-mediated translational restraint, whereas SD supports metabolic restoration and biosynthesis^43,44^.

The mechanisms initiating this response remain unresolved. Volatile anesthetics interact with multiple membrane-embedded targets and substantially alter excitatory and inhibitory synaptic transmission, with effects extending to mechanisms of synaptic plasticity^46^. Such perturbations may contribute to altered cellular homeostasis. Alternatively, volatile anesthetics may perturb lipid membranes and ER homeostasis directly through interactions with embedded protein targets, inducing ISR/UPR activation as a part of a protective proteostatic response.

Importantly, the EEG and transcriptional features following IA were not attributable to systemic stress. Plasma corticosterone levels remained within the expected circadian range after IA, unlike the rise seen after restraint stress, paralleling prior evidence that SD produces only mild, transient hypothalamic-pituitary-adrenal activation^37^. These findings argue that the ISR/UPR program is a first step in cortical cellular recovery rather than a nonspecific stress response.

Previous studies have described overlaps between anesthesia-induced and sleep-related oscillations, but emphasized that their neurophysiologic mechanisms are distinct^6,8,15^. Isoflurane and desflurane, in particular, can partially reduce subsequent NREM slow-wave rebound but often leave REM homeostasis unsatisfied^16,23–25^. We extend these phenomenological studies by showing that IA’s recovery EEG is dominated by a sustained increase in 0-1 Hz power that generalizes across arousal states and persists long after anesthetic clearance. Functional insight is gained by correlated changes of mRNA expression at the molecular level, a key first step impacting translation and protein expression affecting cellular behavior. Prior reports have documented limited, anesthetic-dependent changes in cortical gene expression, including induction of *Fos* and *Egr1*^47,48^. Our ZT6-matched dataset demonstrates that, more broadly, IA and SD drive opposite polarities of stress-gene regulation, with IA engaging proteostatic restraint and SD favoring anabolic permissiveness within the same cortical region and circadian context.

Importantly, there is precedent for anesthetic-induced alterations in neural function persisting after the anesthetic itself has been eliminated. Zurek and colleagues demonstrated that exposure to isoflurane produces a sustained increase in α5-containing GABA_A_-receptor-mediated tonic current and cell-surface receptor expression, providing a mechanistically distinct example of persistent post-anesthetic neuronal alteration^49^. More broadly, persistent alterations in synaptic transmission and plasticity have been proposed as potential substrates for longer-lasting neurocognitive effects of general anesthetics^46^. Our findings extend this concept from receptor- and synapse-level effects to a systems-level recovery phenotype in which altered cortical network activity is accompanied by a coordinated transcriptional response.

The persistence of enhanced ultra-slow power during wake after IA further suggests that behavioural arousal state and slow-oscillation propensity can be dissociated for prolonged periods (longer than two days for up to two weeks), following anesthesia. Clinically, such a bias could contribute to early post-anesthetic grogginess, attentional lapses, or impaired sleep consolidation, or other subtle neurological dysfunction despite behavioral wakefulness, phenomena frequently noted in postoperative cognitive studies^22^, consistent with a period of increased patient vulnerability. Raising the possibility that conventional clinical markers of emergence may precede complete physiological and cellular recovery of the brain. This may be particularly relevant in patients with reduced neurological reserce following prolonged or repeated anesthetic exposure.

At the molecular level, ISR/UPR engagement implies a protective response transiently prioritizing proteostasis over sleep-related, CNS growth and restoration. Conversely, the SD signature indicates anabolic readiness as rebound sleep begins, consistent with restoration of synaptic and metabolic functionality after prolonged wake^30,45,50^. Although causality between the transcriptional and electrophysiological changes remains to be tested, these opposing responses identify testable mechanisms through which anesthesia and physiological sleep may engage fundamentally different recovery programs. Taken together our findings suggest that behavioural emergence from anesthesia should not necessarily be equated with complete biological recovery of the brain.

## Limitations

There are several limitations in the current work. First, a single sex (male) and brain region (FC) were used; however, both sex and region profoundly influence sleep and transcriptional physiology^43,51,52^ so this experiment should be replicated using females and additional brain regions. Next, RNA-seq was obtained at a single recovery time point (ZT6); denser temporal sampling will be required to define the onset and resolution of ISR/UPR engagement. Additionally, IA and SD differ in experiential content (surgical-plane anesthesia versus enforced wakefulness). Although corticosterone data argue against systemic stress during IA, local neuromodulatory environments may diverge and influence molecular responses. Further, causality remains to be established: manipulating ISR/UPR or mTOR nodes during emergence could test whether these molecular programs are drivers or that the observed EEG and behavioral phenotypes are necessarily upstream of the transcriptomic changes.

Finally, these observations and analyses suggest that IA induces a cellular stress response involving ISR/UPR engagement. Although the electrophysiological changes, including ultra-slow oscillatory activity, are reversible over time, we did not assess the temporal dynamics of the associated transcriptional response. In healthy, young, non-compromised mice, the recovery of EEG patterns suggests that these perturbstions may resolve, however, in clinical settings where anesthesia often occurs in the context of compromised (brain) health, the persistence or cumulative impact of such stress responses remains unclear. A more mechanistic understanding of anesthetic-induced cellular stress, and how it relates to exposure duration, may help clarify risk across patient vulnerable states and pathologies.

## Conclusions

Isoflurane anesthesia and sleep represent fundamentally different kinds of unconsciousness. The gene-expression responses induced by IA and SD (each compared to control sleep) are not only distinct but, with regard to proteostasis and growth-related programs, opposite in direction. Together, these findings provide convergent neurophysiologic and molecular evidence that time spent under anesthesia does not “count” as restorative sleep and that a cellular stress response is a defining feature of the post-anesthetic brain.

## Methods

### Animals

All experiments were performed in accordance with institutional guidelines and approved animal care protocols. Male C57BL/6J mice (*Mus musculus*) aged 8–19 weeks at the time of data collection were used for all experiments. Mice were housed under a 12 h light / 12 h dark cycle (lights on at ZT0) with ad libitum access to food and water.

### EEG/EMG Electrode Implantation

For sleep–wake state analysis, mice (*n* = 15) were implanted with EEG and EMG electrodes using standard procedures as previously described^19^ (Bjorness et al., 2016). Briefly, mice were anesthetized with isoflurane and secured in a stereotaxic apparatus. After shaving and cleaning the scalp, a midline incision was made and the skull exposed. Small burr holes were drilled for placement of stainless-steel screw electrodes over frontal cortex (AP +1.7 mm, ML ±1.77 mm), parietal cortex (AP −1.7 mm, ML +2.0 mm), and occipital cortex (AP −5.5 mm, ML −1.5 mm). Paddle-style EMG electrodes (Plastics One) were inserted bilaterally into the dorsal nuchal muscles.

All electrodes were connected to a six-pin pedestal (Plastics One), secured to the skull with dental cement, and the incision was closed with absorbable sutures and topical antibiotic ointment. Postoperative analgesia was provided using buprenorphine and carprofen. Animals were allowed at least 14 days of recovery prior to experimental procedures.

### EEG/EMG Acquisition and Sleep State Scoring

EEG and EMG signals were acquired using a 15LT amplifier system (Natus Neurology; formerly Grass Technologies). Signals were continuously recorded throughout baseline, intervention, and recovery periods.

Sleep–wake state classification was performed using a custom MATLAB (MathWorks) autosorting algorithm based on standard criteria for wake, NREM (slow-wave sleep), and REM sleep^19^. All scoring was blinded for condition (baseline/IA/SD). All scored files were manually reviewed to confirm accuracy and to flag epochs containing artifacts. Data were scored in 10 s epochs; artifact-containing epochs were excluded from spectral analyses but retained for time-in-state calculations. Sleep episodes were defined as beginning after 30 consecutive seconds of a given state and ending after 30 consecutive seconds of a different state.

### Experimental Paradigms: Baseline, Sleep Deprivation, and Anesthesia Acclimation

Mice were acclimated for two weeks in plexiglass cages suspended above a custom treadmill apparatus under a 12 h light/dark cycle, with food and water available ad libitum, as previously described^19,29^ (Bjorness et al., 2016; Bjorness et al., 2020). During acclimation, the treadmill belt was run daily at 3 cm/s for 10 min to habituate animals to movement and facilitate cleaning. Following acclimation, mice were connected to the recording tether via a commutator and allowed an additional week to acclimate to tethered conditions.

### Baseline Recording

Following acclimation, EEG/EMG signals were recorded continuously for 48 h under undisturbed baseline conditions.

### Sleep Deprivation

Sleep deprivation (SD) was performed for 4 h starting at ZT0 by continuous treadmill movement at 3 cm/s. Mice were able to eat and drink during deprivation. Following SD, the treadmill was stopped and EEG/EMG recording continued during recovery.

### Isoflurane Anesthesia

For anesthesia (IA), mice were removed from the treadmill cage at ZT0 and transferred to an anesthesia setup. Anesthesia was induced with 5% isoflurane and maintained for 4 h at 0.5–2% isoflurane, adjusted to maintain a surgical plane. Depth of anesthesia was assessed every 5–10 min by toe pinch and respiratory rate (target 80–100 breaths/min). Animals were maintained on a heating pad and monitored using a non-invasive temperature probe. After 4 h, mice were returned to the treadmill cage and EEG/EMG recording continued throughout recovery.

### Time in State Analysis

Time spent in wake, NREM, and REM sleep was calculated in 1 h bins and expressed as percentage of total recording time. Group comparisons were performed using two-way ANOVA in GraphPad Prism.

### Spectral Power Analysis

Under baseline conditions, individual 10 s epochs were grouped by vigilance state (wake, NREM, REM). Power spectra were computed in 1 Hz bins up to 50 Hz and normalized within each state to total power for that state. Fractional power values were averaged across epochs to generate state-specific spectral distributions. Recovery spectra were normalized to baseline values to compute recovery-to-baseline ratios. Group differences in spectral power distributions were assessed using two-way ANOVA with factors of group and frequency bin, including interaction terms, followed by Šídák-corrected post hoc comparisons where appropriate.

### NREM Episode Analysis

NREM episode length distributions were analyzed during baseline and recovery. Cumulative episode-length distributions were fit for each animal using a single-exponential plateau model Y=Y_M_ -(Y_M_-Y_0_)*exp(-k*x) implemented in GraphPad Prism using least-squares regression without weighting (Y_0_ = 0, Y_M_ = 1). The rate constant *K* reflects the relative prevalence of short NREM episodes, with larger values indicating greater fragmentation. Group comparisons were performed using non-parametric tests as specified in figure legends.

### RNA Isolation and Sequencing

For transcriptomic analyses, mice underwent the same anesthesia or sleep deprivation protocols described above. Following 4 h of anesthesia, animals were returned to their home cage for 2 h to allow anesthetic washout. Animals were sacrificed at ZT6 to match circadian timing with previously published sleep deprivation datasets^53^.

Brains were rapidly extracted, rinsed in ice-cold RNase-free PBS, and dissected on ice. Frontal cortex (Fig. S4) was isolated, snap-frozen on dry ice, and stored at −80°C.

Total RNA was extracted using TRIzol reagent (Thermo Fisher Scientific) and purified with the RNeasy Mini Kit (Qiagen) following manufacturer instructions. RNA quality and concentration were assessed using an Agilent Bioanalyzer High Sensitivity DNA chip. Libraries were randomized to minimize batch effects and sequenced on an Illumina NextSeq 500 platform to generate paired-end, strand-specific 150 bp reads.

RNA-seq Data Processing and Differential Expression Analysis

Raw reads were quality-filtered using FastQC and Trimmomatic. Filtered reads were aligned to the mouse reference genome (mm10) using STAR. Uniquely mapped reads were counted using HTSeq. Read counts were normalized using the RPKM method implemented in edgeR. Genes with RPKM ≥1.0 in all replicates of at least one condition were retained for downstream analysis. (ADD info on filtering for confounders etc)

Differential gene expression was assessed using DESeq2. Genes with an adjusted *p*-value ≤0.05 and absolute log₂ fold change ≥0.3 were considered differentially expressed. Volcano plots and heatmaps were generated to visualize results.

### Corticosterone Measurements

Separate cohorts of male C57BL/6J mice were used for corticosterone measurements. Animals were acclimated for seven days prior to testing. Sleep deprivation was performed for 6 h starting at ZT0 using treadmill movement. Control animals remained undisturbed in treadmill or home cages. At ZT6, animals were sacrificed by isoflurane induction and decapitation, and trunk blood was collected. Serum corticosterone concentrations were measured using a commercial ELISA kit (ALPCO Mouse and Rat Corticosterone ELISA, Cat. #55-CORMS-E01).

## Code and Data availability

All custom code and raw and processed sequencing data generated and used in this study will be made publicly available after publication.

## Acknowledgements

We are grateful for the members of the Sleep and Circadian Rhythm group at UTSW for their continuous support and fruitful discussion, in particular Dr. Shin Yamazaki, PhD, Dr. Yuxiang Liu, PhD, and Dr. Heidi Pak, PhD.

## Author contributions

Conceptualization: MD, SAL, RWG

Methodology: MD, TEB, RWG

Investigation: MD, TEB, TT, RP

Formal analysis: MD, TEB, AK

Resources: AK, GK, RWG

Visualization: MD

Supervision: RWG

Writing – original draft: MD

Writing – review (all authors) & editing: MD, TEB, SAL, RWG

---

**Figure S1.**
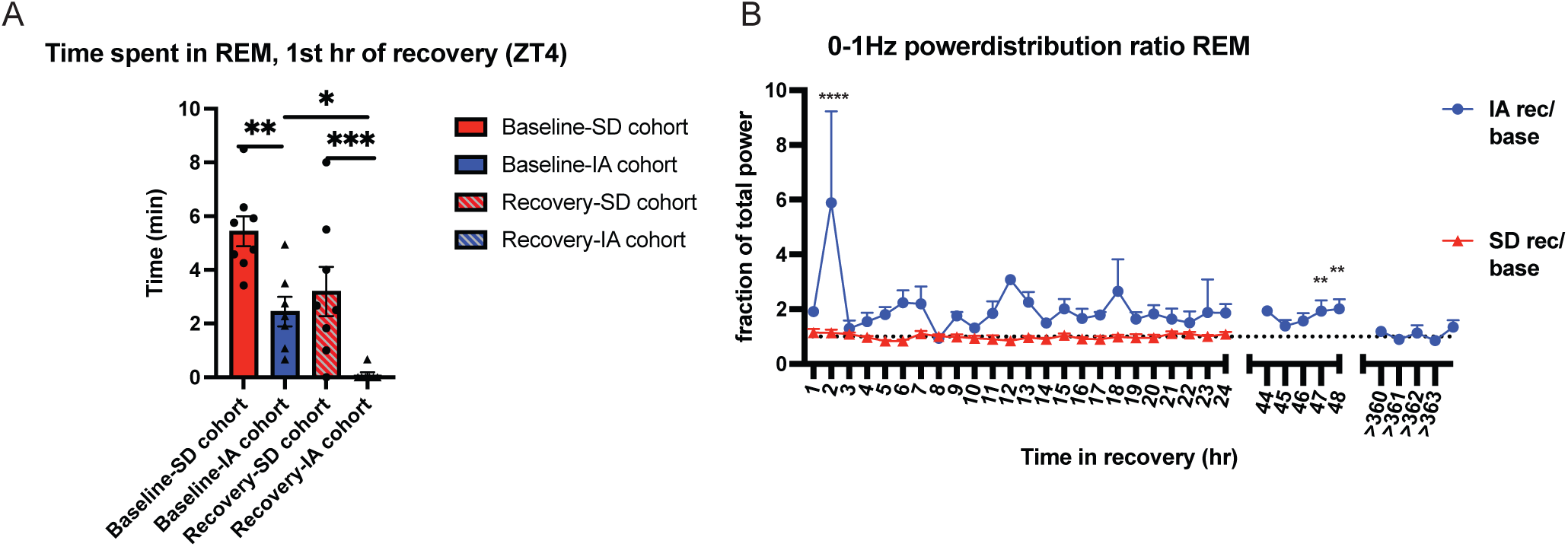
Isoflurane alters REM sleep and ultra-slow power during recovery. (A) Time spent in REM sleep during the first recovery hour (ZT4), expressed relative to the corresponding baseline hour. (B) Recovery-to-baseline ratios for ultra-slow (0-1 Hz) power during REM sleep plotted over two weeks following IA and over the first 24 h following SD. Group comparisons of time spent in REM (A) were performed using a Kruskal-Wallis test with Dunn’s post hoc correction. Group differences in ultra-slow (0-1 Hz) power over time (B) were assessed using a two-way ANOVA with factors of group and time, with Šídák-corrected post hoc comparisons. Data are shown as mean ± SEM. * *p*adj < 0.05, ** *p*adj < 0.01, *** *p*adj < 0.001, **** *p*adj < 0.0001.

**Figure S2.**
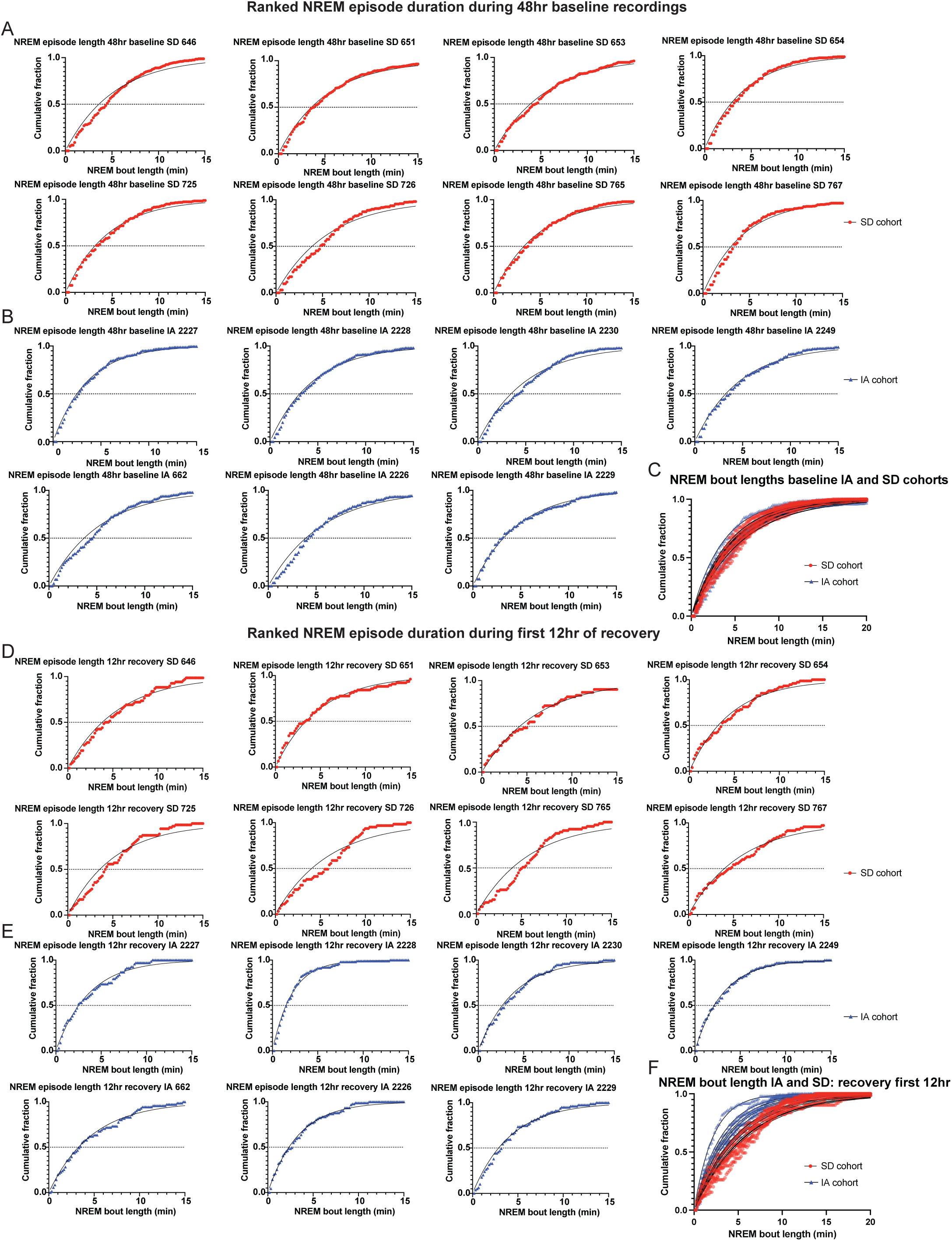
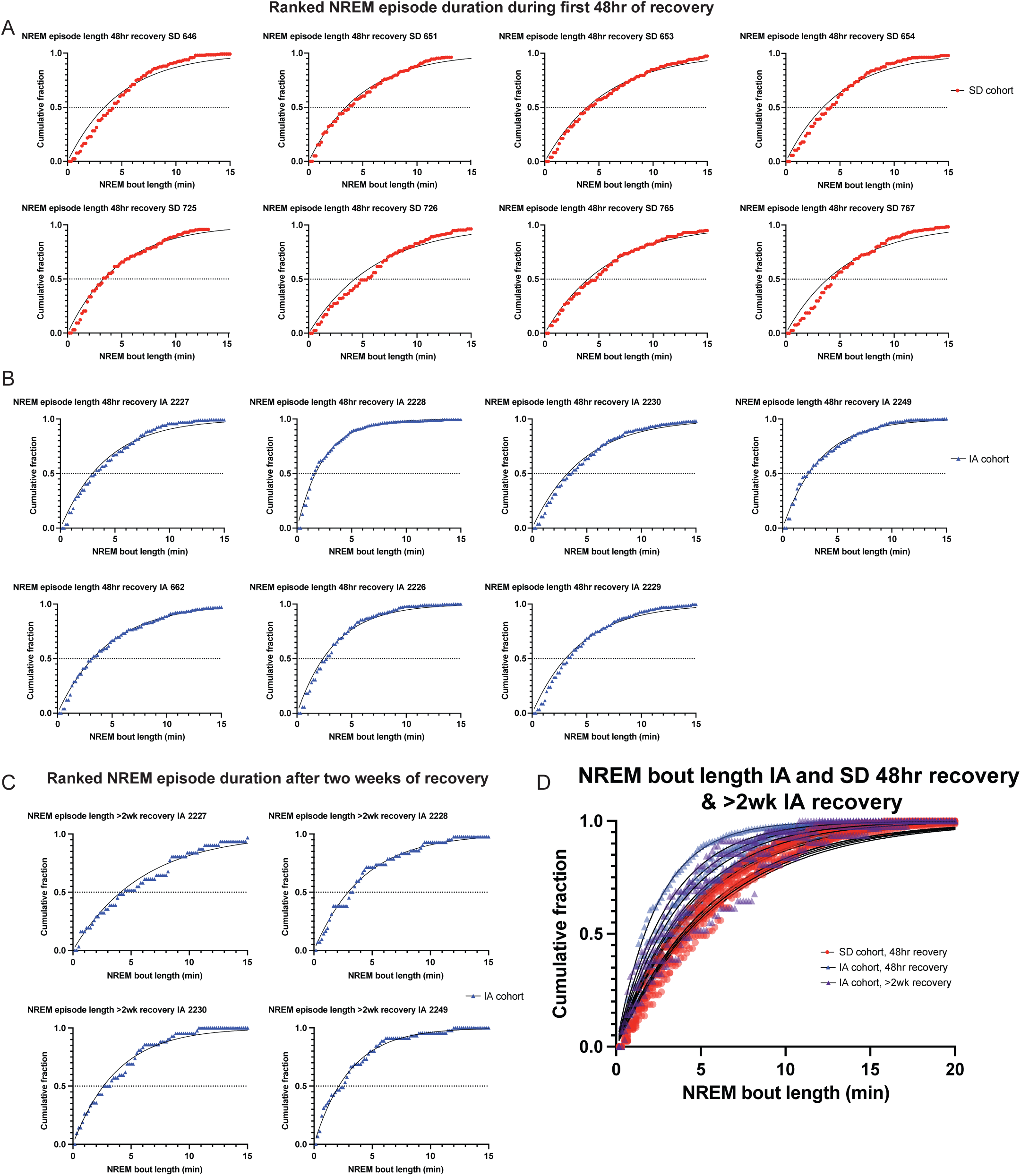
NREM episode length distributions across baseline and recovery. Ranked cumulative NREM episode-length distributions for individual animals. (S2-I-A,B) Baseline recordings for SD (*n* = 8) and IA (*n* = 7) cohorts. (S2-I-C) Overlap of baseline distributions across all animals (data underlying Fig. 1D baseline). (S2-I-D,E) First 12 h of recovery for SD and IA cohorts, respectively. (S2-I-F) Overlap of recovery distributions highlighting cohort separation (data underlying Fig. 1D recovery). (S2-II-A,B) First 48 h of recovery for SD and IA cohorts. (S2-II-C) Recovery distributions >2 weeks post-IA (*n* = 4). (S2-II-D) Overlap of SD and late-recovery IA distributions demonstrating return to baseline architecture.

**Figure S3.**
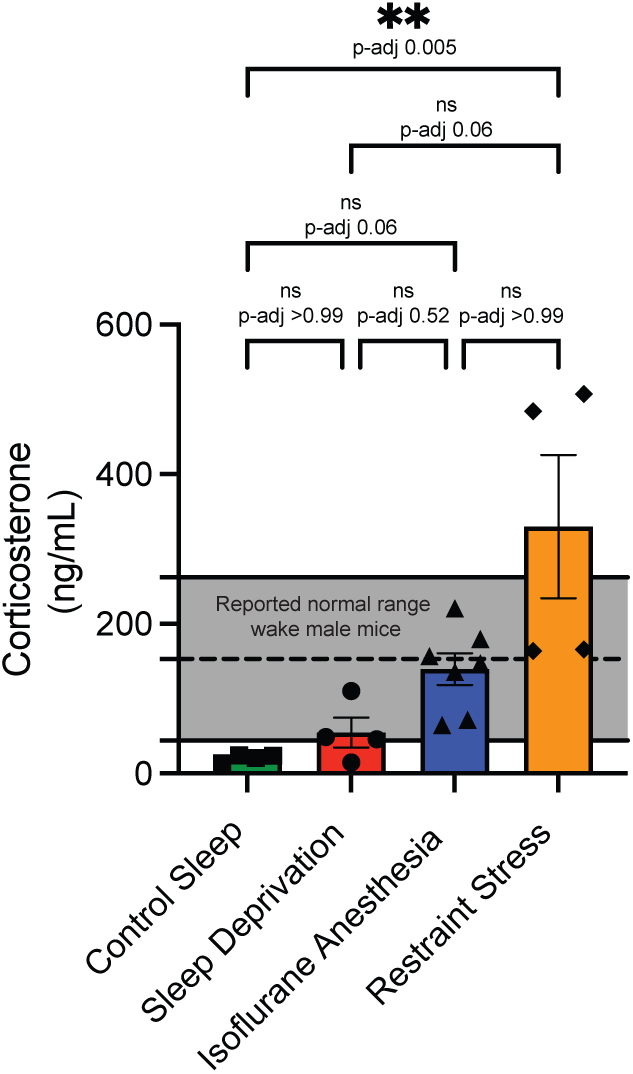
Corticosterone levels following anesthesia and sleep deprivation. Plasma corticosterone concentrations measured by ELISA. Both IA and SD cohorts fall within reported baseline corticosterone ranges for unstressed mice, in contrast to levels observed following moderate restraint stress (*n* = 4).

**Figure S4.**
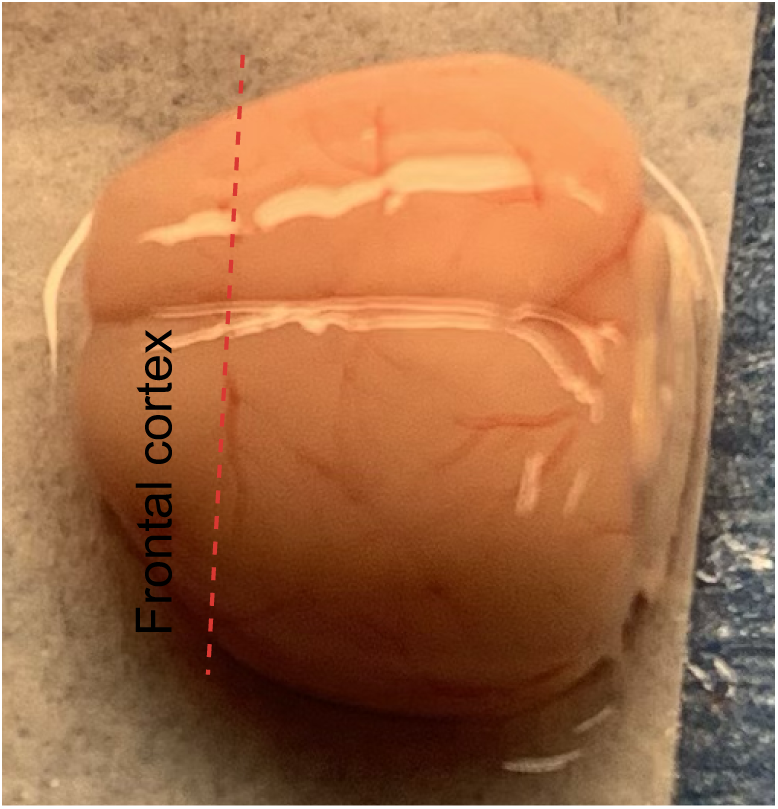
Frontal cortex segment used for bulk RNA sequencing. Cut made at dashed red line, each hemisphere is snap frozen separately. Only one of the hemispheres is sequenced per animal.

